# Decoding of YAP levels and dynamics by pluripotency factors

**DOI:** 10.1101/2022.10.17.512504

**Authors:** Kirstin Meyer, Nicholas C. Lammers, Lukasz J. Bugaj, Hernan G. Garcia, Orion D. Weiner

## Abstract

YAP is a transcriptional regulator that controls pluripotency, germ layer specification, and proliferation. Different subsets of YAP target genes are engaged in each physiological setting, but how YAP selectively regulates different effectors in different contexts is not known. Here we use optogenetics to investigate how the levels and dynamics of YAP activation control its pluripotency effectors Oct4 and Nanog. We observe different thresholds for repression of Oct4 and Nanog, enabling differential control of both genes through YAP levels. Pluripotency factors also decode YAP dynamics. Oct4 preferentially responds to oscillatory YAP inputs that mimic endogenous pulsatile YAP dynamics. Using single-cell live imaging of Oct4 transcription and computational-theoretical analysis of transcriptional regulation, we demonstrate that YAP dynamics are decoded by an adaptive change sensor that modulates Oct4 transcription burst frequency. Our results reveal how the levels and timing of YAP activation enable multiplexing of information transmission for key regulators of cellular differentiation and pluripotency.

## Introduction

Transcription factors play pivotal roles in the regulation of development. They relay information from the cellular environment to the gene regulatory networks that control cell fate. A striking feature of these transcriptional networks is the use of a relatively small set of transcription factors to control large arrays of genes during development, with the same transcription factors often regulating different sets of genes in different contexts (Heng et al., 2020; Lavoie et al., 2020; Nandagopal et al., 2018). When a transcription factor is activated, how do cells choose among multiple downstream responses? Addressing this question is not only fundamental for developmental biology but will also be essential for manipulating cell fate for tissue engineering.

YAP (yes-associated protein), the main effector of the Hippo pathway, is a functionally pleiotropic transcriptional regulator. It integrates inputs from cell mechanics (Aragona et al., 2013; Dupont et al., 2011), cell polarity (Yang et al., 2015), and cell metabolism (Anakk et al., 2013; Yu et al., 2012) to control the effectors of pluripotency (Lian et al., 2010; Qin et al., 2016), germ layer specification (meso-/endo-/ectoderm, (Chung et al., 2016; Stronati et al., 2022)) and proliferation (Wu et al., 2003; Zhao et al., 2007). While extensive efforts have identified the signaling modules that act upstream and downstream of YAP, we lack an operational framework for understanding how YAP controls specific gene regulatory programs and cellular decisions.

In many biological contexts, signaling levels and dynamics are used to specify cellular responses (Hao and O’Shea, 2011; Purvis and Lahav, 2013). For example, different concentrations of morphogens direct distinct gene regulatory programs, enabling the conversion of continuous morphogen gradients into switch-like boundaries of cell fate (Jiang and Levine, 1993; Melen et al., 2005). Alternatively, the temporal dynamics of signaling inputs (such as the duration, frequency and amplitude) can be used to specify appropriate cellular behavior. For example, ERK and p53 dynamically inform downstream target genes about the identity and magnitude of upstream inputs, respectively. ERK target genes act as persistence detectors to control whether cells proliferate or differentiate (Heasley and Johnson, 1992; Toettcher et al., 2013; Traverse et al., 1992). p53 pulse frequency conveys the magnitude of DNA damage; this enables cells to decide between DNA repair and cell death (Batchelor et al., 2011; Lahav et al., 2004). Despite fluctuations of YAP levels in developmental contexts (Franklin et al., 2020), it is unclear if cells employ a similar dynamic or concentration-dependent encoding strategy to link patterns of YAP activation to appropriate downstream effectors (Fig. 1A).

**Fig. 1.**
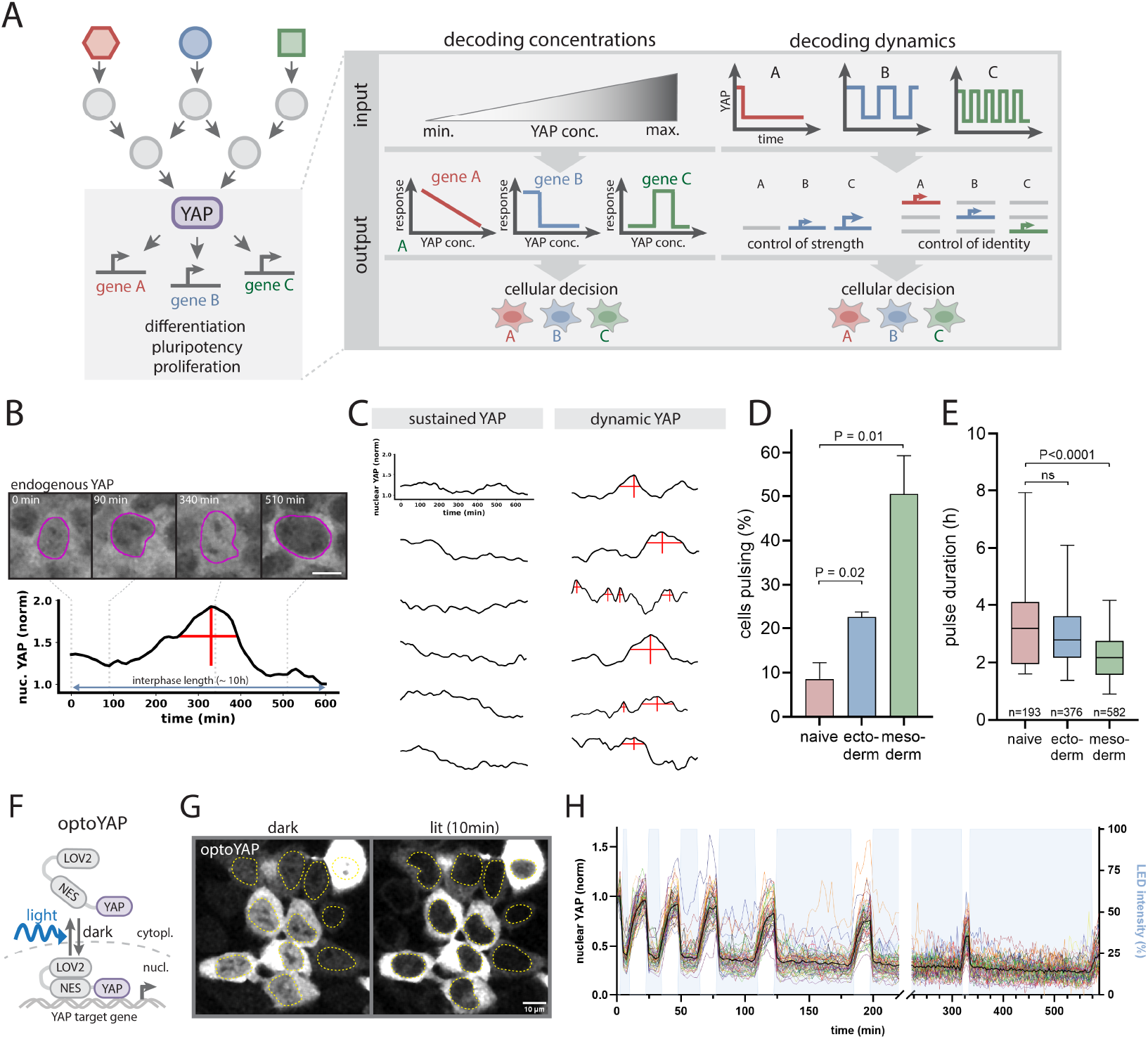
Light-gated control of nuclear YAP levels mimics endogenous YAP dynamics. A) Hourglass-shaped topology of the YAP signaling network. YAP concentrations and dynamics could control downstream effector strength and/or identity. B) Pulsatile nuclear dynamics of endogenously tagged SNAP-YAP in interphase mESCs at 36h post directed mesoderm induction. Scale bar: 10 μm. C) Representative single-cell quantitation of nuclear YAP levels in the same culture condition as (B). Traces are classified as sustained or dynamic based on an automated peak detection algorithm. Peak width and height are indicated by horizontal and vertical red lines. D, E) Quantification of the percentage of cells exhibiting YAP pulses (D) and the average YAP pulse duration (E) in naive mESCs versus 36h directed induction along the ectoderm or mesoderm lineages. Cells were classified by the peak detection strategy shown in (C). Shown are mean +/-SEM, N=3 (C) and the Box and Whiskers with median + 5-95 Percentile, n as indicated, N=3. (D). p values from unpaired Student’s t test. F) Optogenetic strategy to control nuclear YAP levels by light using the LEXY-tag. LEXY leverages the engineered LOV2 protein domain that unfolds a nuclear export sequence (NES) upon blue light illumination, causing reversible nuclear optoYAP export. G) Representative images of mESCs expressing optoYAP in the dark (pre-activation) and following 10 min light illumination (post-activation). Nuclei are outlined in yellow. Scale bar: 10 μm. H) Quantification of nuclear optoYAP levels from microscopy time course demonstrating light-gated induction of consecutive nuclear YAP import/export cycles ranging from the minute to hour timescale. Illumination phases are indicated by blue shading.

Here we investigate how the levels and timing of YAP control downstream target activation in mouse embryonic stem cells. This analysis necessitated more sophisticated temporal control of nuclear YAP levels and dynamics than can be achieved through classical genetic perturbation strategies. For this purpose, we combined an optogenetic system to dynamically control nuclear YAP levels with live imaging of downstream target gene transcription to reveal the input-output logic of YAP signaling. We demonstrate the differential control of the pluripotency factors Oct4 and Nanog through YAP levels and dynamics. Oct4 and Nanog display distinct thresholds for inhibition by YAP, enabling differential control of these genes. By applying light-gated oscillatory YAP dynamics we further reveal a dynamic decoding capacity of Oct4, which acts as an adaptive change sensor that optimally engages at specific YAP frequencies mimicking those found in the endogenous system. Our work reveals how the levels and timing of YAP activation enable multiplexing of information transmission in development. This work could establish a foundation for precise control of cell fate through synthetic control of YAP dynamics.

## Results

### mESCs induce YAP dynamics during differentiation

YAP is a primary determinant of cell fate during early development (Chung et al., 2016; Stronati et al., 2022). To assess if YAP levels or dynamics could control mESCs cell fate decisions, we first analyzed native YAP dynamics in mESCs. For this purpose, we generated an endogenous SNAP-YAP reporter mESC line and monitored YAP levels at 1.5d post differentiation. This time point was chosen because it is a generally permissive time window for differentiation cues in mESCs (Semrau et al., 2017). Live-cell imaging of the meso- and ectodermal lineages (differentiation efficiency > 50%, Fig. S1A-C) revealed temporal fluctuations of nuclear YAP levels (Fig. 1B, C, Movie S1). While a small proportion (<10%) of naive cells show nuclear YAP fluctuations (Fig.1D), 23%-50% of mESCs differentiating into the ecto- and mesodermal lineages exhibit discrete YAP pulses (Fig.1B-D), with an average 1.5-fold change in amplitude (Fig. S1D) that last on average 2.3-3h (Fig. 1E). The observation that mESCs exhibit YAP dynamics during the onset of differentiation is consistent with the hypothesis that cells employ a YAP concentration-dependent or temporal decoding strategy for controlling cell fate.

### Inducible and light-gated control of nuclear YAP levels in mESCs

To investigate whether the levels or dynamics of YAP play an instructive role in downstream effector activation, we adopted a previously reported optogenetic tool termed iLEXYi (Fig. 1F; (Kögler et al., 2021; Niopek et al., 2016) to control nuclear YAP levels by light. iLEXYi is based on the AsLOV2 domain (Christie et al., 1999; Wu et al., 2009) that exposes a nuclear export sequence upon blue light illumination, resulting in reversible nuclear export. We fused iLEXYi to fluorescently-tagged YAP (iLEXYi-SNAP-YAP termed optoYAP) and expressed it in a YAP KO background (Fig. S1E) under a doxycycline-inducible promoter. This enabled us to control both long-term titration of steady-state YAP levels by doxycycline as well as acute modulation of nuclear YAP dynamics by light. Doxycycline dose-response analysis demonstrated our ability to achieve a wide range of steady-state optoYAP expression levels that bracket the endogenous YAP levels of WT mESCs (Fig. S1F). With our optogenetic system, we can achieve rapid and reversible nuclear-cytoplasmic shuttling on the minute timescale (export 5min, import 15min, Fig.1G,H, Movie S2) with a maximum nuclear YAP depletion of 60% (Fig. S1G). Illumination with different light durations enables pulse width modulation ranging from minutes to hours over extended time periods (10h; Fig.1H) that mimic the amplitude and duration of the temporal changes observed in endogenous contexts (see Fig. 1E, Fig. S1D). We leveraged our optoYAP tool to investigate concentration- and time-dependent responses of YAP effectors. In addition to probing YAP levels and dynamics that mimic those observed in the endogenous system, we also investigated a larger parameter space to infer general signaling principles. First, we analyze how the levels of YAP control gene activation. Next, we ask how the temporal dynamics of YAP control gene activation. Finally, we investigate how downstream effectors decode the levels and timing of YAP activation.

### Steady-state YAP levels differentially control pluripotency factors Oct4 and Nanog

To probe the logic of YAP decoding at the molecular level, we focused on the YAP targets Oct4 and Nanog (Fig. 2A top panel), which play critical roles in pluripotency maintenance and differentiation (Loh et al., 2006; Thomson et al., 2011). We titrated steady-state YAP levels and measured gene doseresponse (Fig. 2A bottom panel) through immunofluorescent (IF) staining of Oct4 and Nanog (Fig. 2B). To cover a wide range of YAP levels, we made use of the expression heterogeneity of the doxycycline inducible system and analyzed cells expressing SNAP-YAP at 2 days following spontaneous differentiation. Here and for all other physiological measurements, we grew cells on PDMS substrates of physiologically relevant stiffness (64kPa) to ensure a defined permissive mechanical environment (see Methods for details), as previously established for YAP’s role in cellular reprogramming (Chang et al., 2018). Our dose-response experiments identify YAP as a repressor of both Oct4 and Nanog (Fig. 2B,C). Notably, the proteins show different thresholds for inhibition by YAP with IC50s (Fig. 2C, dotted lines) that establish a window for the differential control of Oct4 and Nanog. Within this window, there are levels of YAP that significantly repress Nanog and not Oct4. Above and below this differential control window, YAP acts jointly on Nanog and Oct to permit expression (low YAP) or induce repression (high YAP) of both genes. Naïve mESCs are less sensitive to YAP levels with a significant shift of the Oct4 and Nanog IC50s as compared to differentiating cells (Fig. S2A), indicating that YAP signaling competence changes during pluripotency exit. Importantly, the observed dose-response behavior is comparable between cells expressing SNAP-YAP and optoYAP (Fig. S2B), verifying the functionality of our optogenetic tool. Our results demonstrate differential control of gene activation through steady-state YAP levels, analogous to the role of morphogens in determining cell fate in a concentration-dependent manner. We next sought to investigate how the dynamics of YAP control gene activation using our optogenetic approach.

**Fig. 2.**
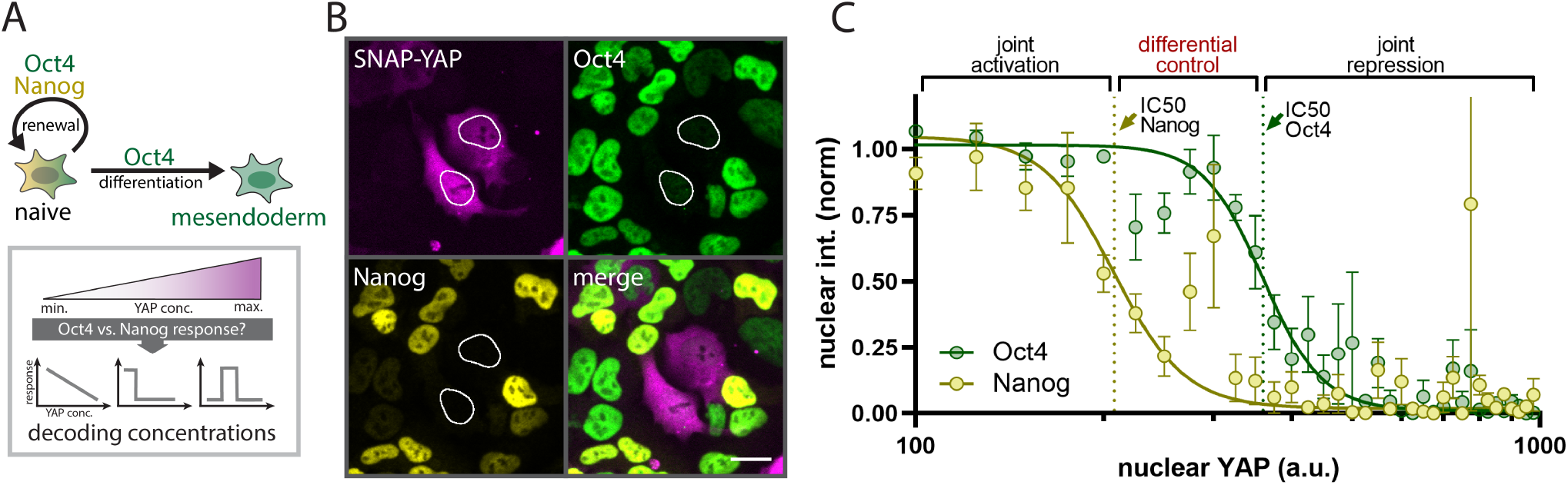
Differential control of Oct4 and Nanog through steady-state YAP concentrations. A) Top: The YAP target genes Oct4 and Nanog work together to regulate naive mESC renewal, while Oct4 alone promotes differentiation into the mesendoderm fate. Bottom: Probing the dose response of Oct4 and Nanog accumulation as a function of steady-state YAP concentrations. B) SNAP-tag and immunofluorescent staining for SNAP-YAP, Oct4 and Nanog at 48h post undirected differentiation start shows that YAP is a repressor of Oct4 and Nanog. Nuclei with high SNAP-YAP expression are outlined in white. Scale bar: 20 μm. C) Hill curve fit of nuclear Nanog and Oct4 levels as a function of nuclear YAP concentrations reveals differential sensitivity of Oct4 and Nanog to YAP levels through offset repression thresholds (IC50s) that establish three distinct regimes for joint activation of Oct4 and Nanog (low YAP levels), joint repression of Oct4 and Nanog (high YAP levels) and preferential inhibition of Nanog and versus Oct4 (intermediate YAP levels). Dashed lines indicate IC50s. Shown are mean +/-SEM, N=4.

### The pluripotency factors Oct4 and Nanog decode YAP dynamics

To test how the timing of YAP activation modulates Oct4 and Nanog regulation, we used our optoYAP tool to dynamically control the concentration of YAP in the nucleus. We induced light-gated oscillatory YAP dynamics with constant export duration (4h) but varying recovery (import) periods for a total of 12h during the onset of differentiation (24h-36h post pluripotency exit; Fig. 3A). We investigated YAP pulse durations ranging from 15min to 4h that bracket the range of temporal pulse features observed in the endogenous system (see Fig. 1E). IF staining and single-cell quantification of Oct4 and Nanog protein levels reveal that Oct4 and Nanog are more potently induced by dynamic oscillatory YAP inputs than they are by chronic YAP inputs (Fig. 3B-D). While chronic YAP export caused a moderate increase of Oct4 and Nanog (Oct4: 1.4-fold; Nanog: 1.1-fold, Fig. 3 C,D), oscillatory light inputs induce significantly higher protein levels reaching a maximum induction upon illumination with a 4h export/1h recovery duty cycle (Oct4: 2.2-fold; Nanog: 1.3-fold, Fig. 3C,D). Interestingly, very long import cycles (4h) are less effective than shorter ones (e.g. 1h) demonstrating a dynamic filtering capacity. In conjunction with our steady-state experiments, our data show that both the levels and timing of YAP activation differentially engage the YAP downstream targets Oct4 and Nanog through different regulatory modes. While steady-state levels differentially control Oct4 and Nanog within a fixed concentration regime, dynamic YAP inputs jointly induce Oct4 and Nanog, although with different magnitudes (Oct4: 2.2-fold vs Nanog 1.3-fold induction). Given the importance of transcription factor ratios, for example during cellular reprogramming, these difference in magnitude could establish differential control regimes similar to the threshold-dependent decoupling of Oct4 and Nanog expression under steady-state YAP concentrations. Together, these identified regulatory modes could provide a means for the complex regulatory requirements of Oct4 and Nanog: pluripotency maintenance requires overlapping high Oct4 and Nanog expression (achievable at low YAP), while lineage commitment requires mutually exclusive control (low Nanog and high Oct4, achievable at intermediate YAP or dynamic YAP inputs) or overlapping low expression (low Nanog and low Oct4, achievable at high YAP).

**Fig. 3.**
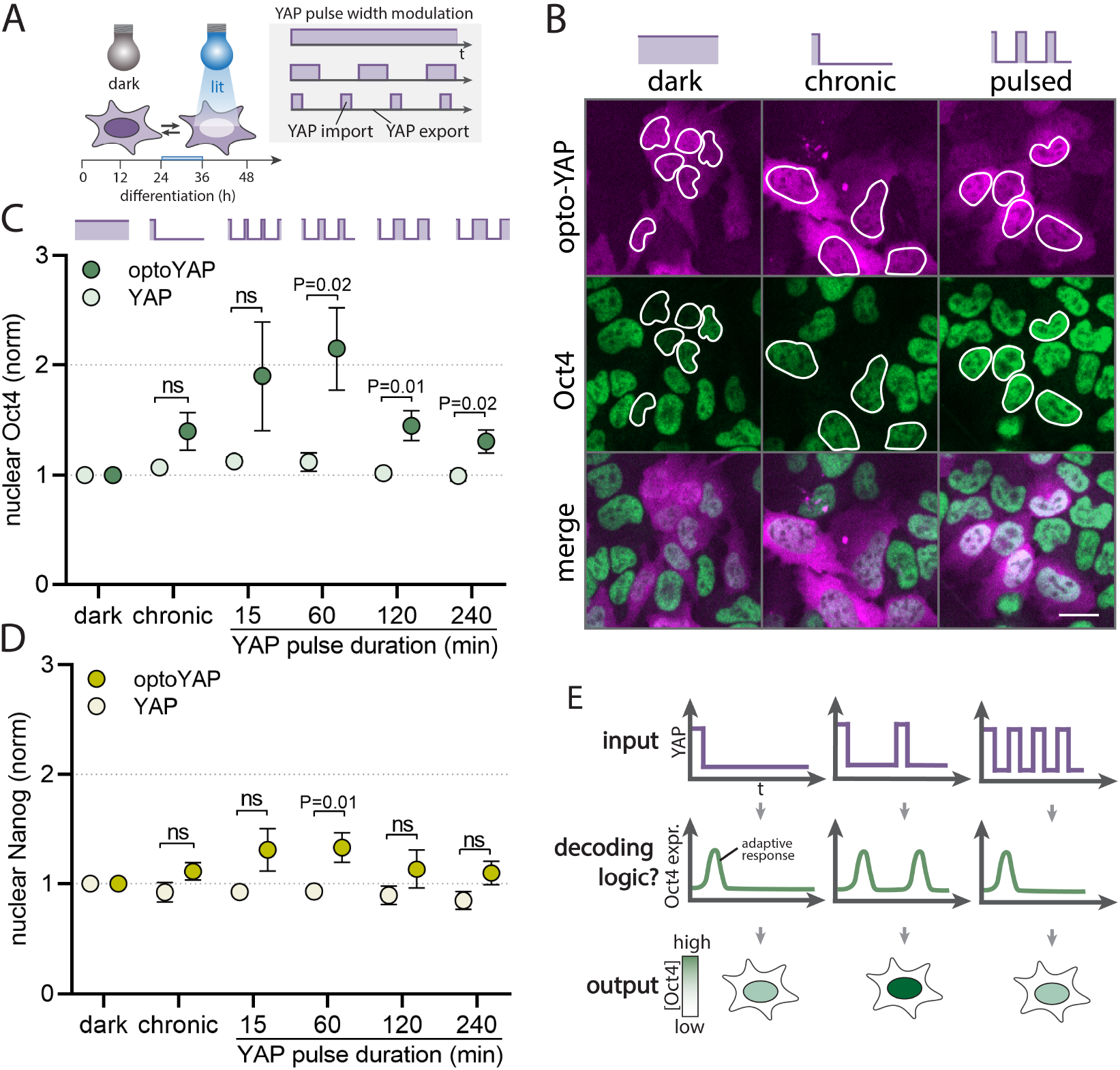
Oct4 and Nanog respond to YAP dynamics. A) To probe whether YAP targets are sensitive to dynamic YAP inputs, ES cells expressing optoYAP were exposed to light patterns with different pulse widths during pluripotency exit. B) IF staining for Oct4 in optoYAP mESCs upon illumination with chronic or pulsed light inputs demonstrates induction of Oct4 upon illumination as compared to the dark control, with higher Oct4 levels in the oscillatory than chronic YAP export condition. Scale bar: 20 μm. C, D) Quantification of nuclear Oct4 (C) and Nanog (D) levels from IF staining as shown in (B). Cells were subjected to light pulses inducing constant YAP export durations of 4h and varying import duration ranging from 15min to 4h as indicated. Pulse frequencies with specific YAP export/import cycles (4h export, 60min import) significantly upregulate Oct4 and Nanog as compared to the non-light responsive YAP control or compared to chronic YAP export. Shown are mean+/-SEM, N=7, p values from unpaired Student’s t test. E) Possible decoding logic of dynamic YAP inputs through adaptation. For an adaptive system, continuous YAP export would transiently activate YAP effectors (left panel). In contrast, pulsed YAP inputs would induce sequential adaptive YAP responses; this would result in higher total Oct4 induction (center panel) than is seen for chronic YAP export (left panel). The adaptive system gives an optimal output at a specific pulse frequency input (center) compared to faster (right panel) or slower (not shown) frequencies.

Our data shows optimal activation of YAP targets at a particular frequency of YAP activation. What is the basis of this temporal decoding? One potential mode of preferentially responding to a given signaling dynamics is the adaptive circuit found in other systems such as chemotaxis (Barkai and Leibler, 1997), osmoregulation (Muzzey et al., 2009), sensory systems (Kurahashi and Menini, 1997; Pugh et al., 1999), and for other transcription factors (Nelson, 2004; Ryu et al., 2016). In response to an acute chronic input, adaptive signaling circuits transiently respond to the change in signal input but then reset to the initial baseline under sustained activation of the input (Fig.3E, left). Adaptive systems require resetting between rounds of activation and only produce a single pulse for oscillatory inputs that are too closely spaced in time (Fig. 3E right panel). As a result, adaptive systems generate optimal responses at a specific input frequency that matches the resetting time (Fig. 3E). We next investigated whether Oct4 uses this adaptive strategy to decode the temporal dynamics of YAP.

### Oct4 acts as an adaptive change sensor of YAP

To investigate how Oct4 expression responds to acute changes in YAP, we leveraged the MS2 system to visualize real-time transcription in individual living cells (Bertrand et al., 1998). The MS2 system is based on the integration of a repetitive MS2 DNA array into the endogenous gene locus (Fig. 4A). Transcription of the array generates hairpin structures at the RNA level that are detectable through recruitment of a Halo-tagged coat protein. Local spots of labeled RNA are visible at the transcription site and provide a measure of transcriptional activity of the gene locus. We integrated the MS2 array into the endogenous Oct4 locus in WT and YAP KO background mESCs and observed sporadic bursts of Oct4 RNA production (Fig. 5B) that are characteristic features of transcriptional activation. The MS2 spots colocalized with the Oct4 locus by DNA FISH (Fig. S3A). To verify that YAP is controlling Oct4 at the gene regulatory level, as expected for a transcriptional regulator, we compared the Oct4-MS2 reporter in YAP KO to WT cells at 2d following pluripotency exit. While WT cells show moderate transcriptional activity (on average 18+/-4.2% (mean+/SEM) of the cells are transcriptionally active), depletion of YAP strongly potentiates Oct4 expression, with an average of 62+/-4.6% (mean+/-SEM) cells bursting (Fig. S3B,C; Movie S3). This confirms the role of YAP as a transcriptional repressor of Oct4. To analyze how Oct4 transcriptional activity responds to steady-state YAP levels and dynamics, we used the Oct4-MS2 reporter to monitor transcriptional activity before and after acute light-gated nuclear export. Mapping the dark (pre-activation) phase single-cell Oct4-MS2 activity to steady-state nuclear optoYAP levels, we found a similar concentration-dependent repressive effect (Fig. 4C) as observed at the protein level (see Fig. 2C), demonstrating that the steady-state control of Oct4 is implemented at the level of transcription. Co-staining of optoYAP cells and WT mESCs for YAP reveals that the dose-responsive regime of the optoYAP tool falls into the endogenous YAP concentration range of WT cells (Fig. 4C, gray shading), indicating that our optoYAP tool operates in a physiological range.

**Fig. 4.**
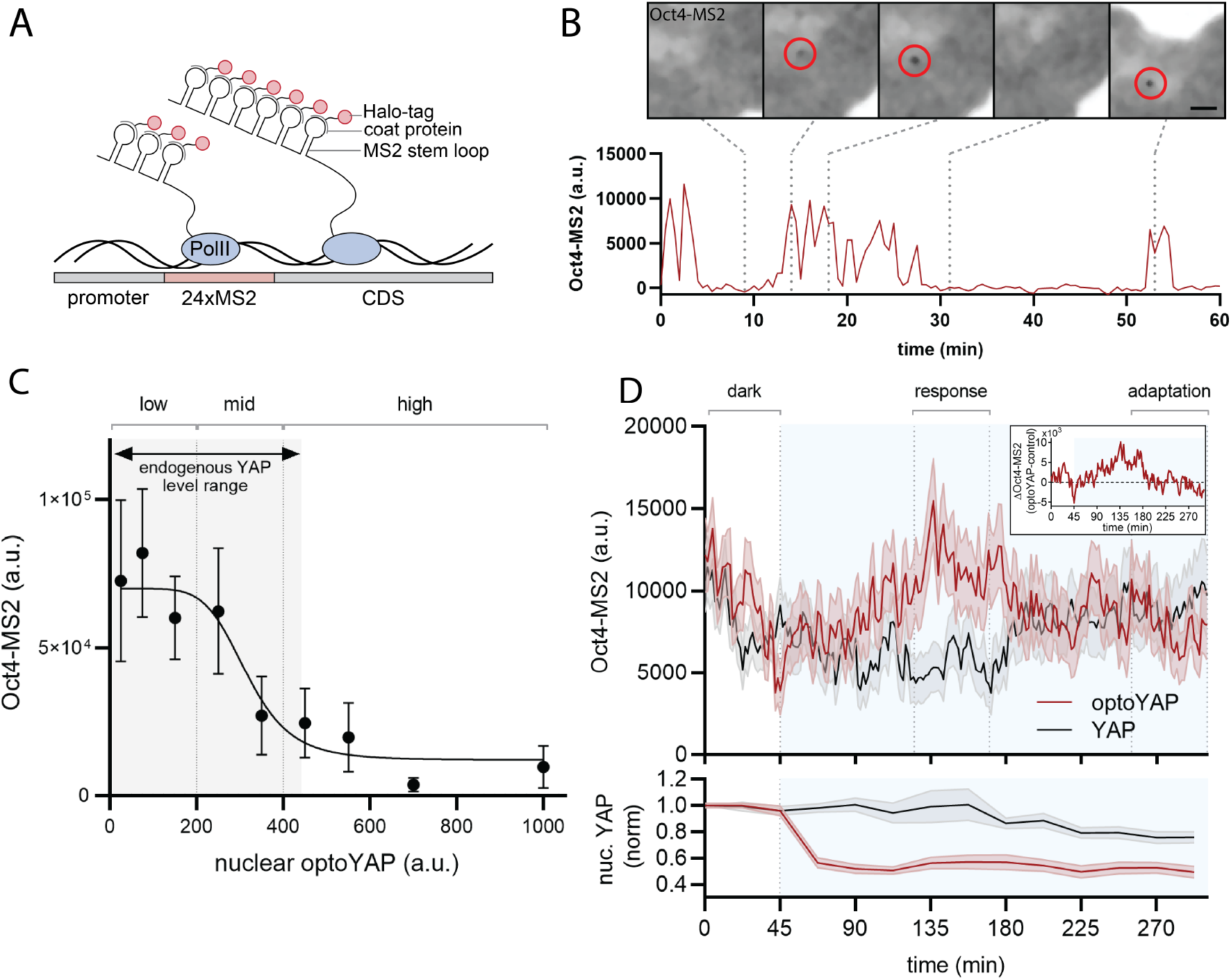
Oct4 acts as adaptive change sensor of YAP levels. A) MS2 system for visualization of transcription in single living cells. Transcription of the 24xMS2 DNA array generates RNA stem loops that are detected through their recruitment of Halo-tagged coat proteins and visible as fluorescent spots at the transcription site (see B). CDS: Coding sequence. B) Oct4-MS2 live-imaging reporter in WT mESCs shows transcriptional bursting. Example microscopy data for indicated time points (dashed line) of the time course. Scale bar: 2.5 μm. C) Oct4-MS2 signal as a function of nuclear optoYAP levels reveals that YAP acts as a repressor of Oct4 transcription. Min-max range of endogenous YAP levels measured in WT cells is indicated by gray shading. Shown are mean +/-SEM., N=15. D) Top: Oct4-MS2 transcriptional activity upon light-gated nuclear YAP export in optoYAP mESCs (red curve) vs. non-light responsive YAP control cells (gray curve). Difference between mean of optoYAP and YAP curve is shown as inset on the top right, indicative of transient adaptive YAP target activation in response to sustained export of YAP from the nucleus. Bottom: Nuclear YAP levels in optoYAP (red) and YAP (gray) mESCs simultaneously imaged for Oct4-MS2 shown in the top panel. Illumination phase is indicated by blue shading. Shown are mean +/-SEM, n > 48 per condition, N =12.

Next, we analyzed the Oct4-MS2 behavior after light-gated YAP export from the nucleus. Considering only optoYAP cells with expression levels in the endogenous range (see Fig. 4C), we observe an adaptive response of Oct4 transcriptional activity, with a transient peak of activation followed by return to baseline levels (Fig. 4D, top panel; Movie S4) in the presence of sustained YAP export (Fig. 4D, bottom panel). The Oct4 MS2 response exhibits an approx. 45min delay relative to the time of initial illumination and lasts for about 1.5h before resetting to the baseline. This response is different from a concentration-dependent mode of regulation where we would expect Oct4 to be persistently activated following sustained YAP export from the nucleus. The time scale of the Oct4 response (45min delay to light onset and 1.5h duration of Oct4 expression) is consistent with our observations at the protein level; these respond to light pulses of 4h pulse width to efficiently increase Oct4 protein levels. The 45min initial delay and long response time of Oct4 to acute YAP changes suggests that pulses with significantly shorter width and reset time should be poor at inducing Oct4. To test this, we compared Oct4 protein levels between cells exposed to chronic light, slow oscillations (4hON/1hOFF) and a very fast 15min ON/OFF duty cycle (Fig. S3F). The results confirm our hypothesis of an optimal frequency of YAP activation, demonstrating that fast oscillations cannot induce Oct4 levels as potent as slower (4h ON, 1h OFF) oscillations. Our results reveal that Oct4 employs an adaption-based decoding mechanism of YAP dynamics with a characteristic response time on the hour time scale that is reminiscent of the dynamics observed for endogenous YAP during directed differentiation.

### YAP regulates Oct4 transcription through modulation of burst frequency

Our results reveal two different YAP decoding modes that are implemented at the gene regulatory level: steady-state YAP levels differentially control Oct4 and Nanog. And YAP dynamics differentially control the magnitude of Oct4 and Nanog induction. What is the underlying gene regulatory logic that establishes these different examples of YAP decoding? As shown in our single cell Oct4-MS2 traces (see Fig. 4B), transcription occurs as sporadic bursts of nascent RNA synthesis. These bursts are shaped by the transcription cycle in which gene promoters switch from an OFF to an ON state to initiate polymerases. To shape gene expression, transcriptional regulators such as YAP can interface with any steps in this cycle to modulate the total RNA output by control of burst initiation, duration, and amplitude. For example, YAP could act through chromatin modification or recruitment of polymerases to switch gene loci into an ON state and initiate transcription. The different YAP signaling modes observed (Nanog vs Oct4 and steady state vs dynamic) could either be established through differences in how YAP interfaces with the transcriptional cycle or through different YAP sensitivities of individual transcription regulatory nodes. Biochemical snapshots of YAP interaction partners and chromatin state are insufficient to reveal the regulatory entry points shaping the dynamic process of YAP-mediated transcription. Here we investigate the decoding logic of YAP concentrations and dynamics by applying a previously-reported theoretical approach to infer promoter states from our experimental transcription traces.

RNA polymerases take minutes to read through a gene locus. The MS2 signal recorded by our live imaging approach therefore only reflects an integrated readout of transcriptional activity over the elongation time. To deconvolve the instantaneous promoter states from the MS2 signal, we use a previously-reported compound-state Hidden Markov Model (Lammers et al., 2020) that describes transcriptional bursting as a stochastic process in which the promoter switches between an ON and OFF state with kon and koff rates, and initiates polymerases at rate r when in the ON state (Fig. 5A top panel). The rate constants define transcriptional features and enable us to infer burst frequency (kon), duration (koff) and amplitude (r; Fig. 5A bottom panel) to compare principles of gene regulation both between loci and under different YAP inputs (concentration vs dynamics). First, we investigated the molecular logic of how steady-state YAP levels mediated gene repression of Oct4 and Nanog. To this end, we generated a Nanog-MS2 knock-in line in the WT and YAP KO backgrounds to compare Oct4 with Nanog burst features in YAP-depleted mESCs to those of WT cells. Despite their differential sensitivity to YAP levels (see Fig. 2C), inference analysis reveals a surprisingly similar YAP regulatory logic for both genes that acts through modulation of burst initiation. YAP depletion increases burst frequency of both genes (Oct4: 13-fold, P<0.002; Nanog: 2.6-fold, P=0.12, not significant) as compared to WT cells. The Oct4 locus additionally shows a 4.4-fold increase in burst amplitude that is different from the regulation of Nanog but less pronounced than the increase in frequency. Together, these similar regulatory logics suggest that the observed differential control of Oct4 and Nanog through steady-state YAP levels is based on similar decoding mechanisms but different sensitivities, for example through differences in the number or affinity of binding sites. Next, we compared the transcriptional burst modulation between our Oct4 steady-state dose response (Fig. 4C) and dynamic adaptive response (see Fig. 4D). To this end, we classify cells into low, middle and high YAP expressors for the dose-response experiments (see Fig. 4C) and into dark or early and late lit phase for the temporal analysis of the adaptive response (see Fig. D). Interestingly, and consistent with the YAP knockout phenotype, both YAP decoding modes (steady-state and dynamic YAP) are established through the same transcription regulatory logic by modulation of burst frequency. While burst frequency gradually decreases with higher YAP levels, the adaptive Oct4 response is based on a transient 3.5-fold increase in burst initiation rate as compared to the control. Consistent with the adaptive nature of the MS2 signal, burst frequency almost fully resets to the dark-state baseline after 4h. As for the knockout phenotype, both the steady-state and the adaptive response are accompanied by a small increase in burst amplitude, suggesting that YAP may act on more than one regulatory node for the Oct4 locus. Together, the results demonstrate that the adaptive change sensor and dose response module makes use of the same transcription regulatory machinery, suggesting that differences in the interpretation of steady-state and dynamic YAP inputs must be established through other regulatory feedback loops, possibly at the level of YAP itself.

**Fig. 5.**
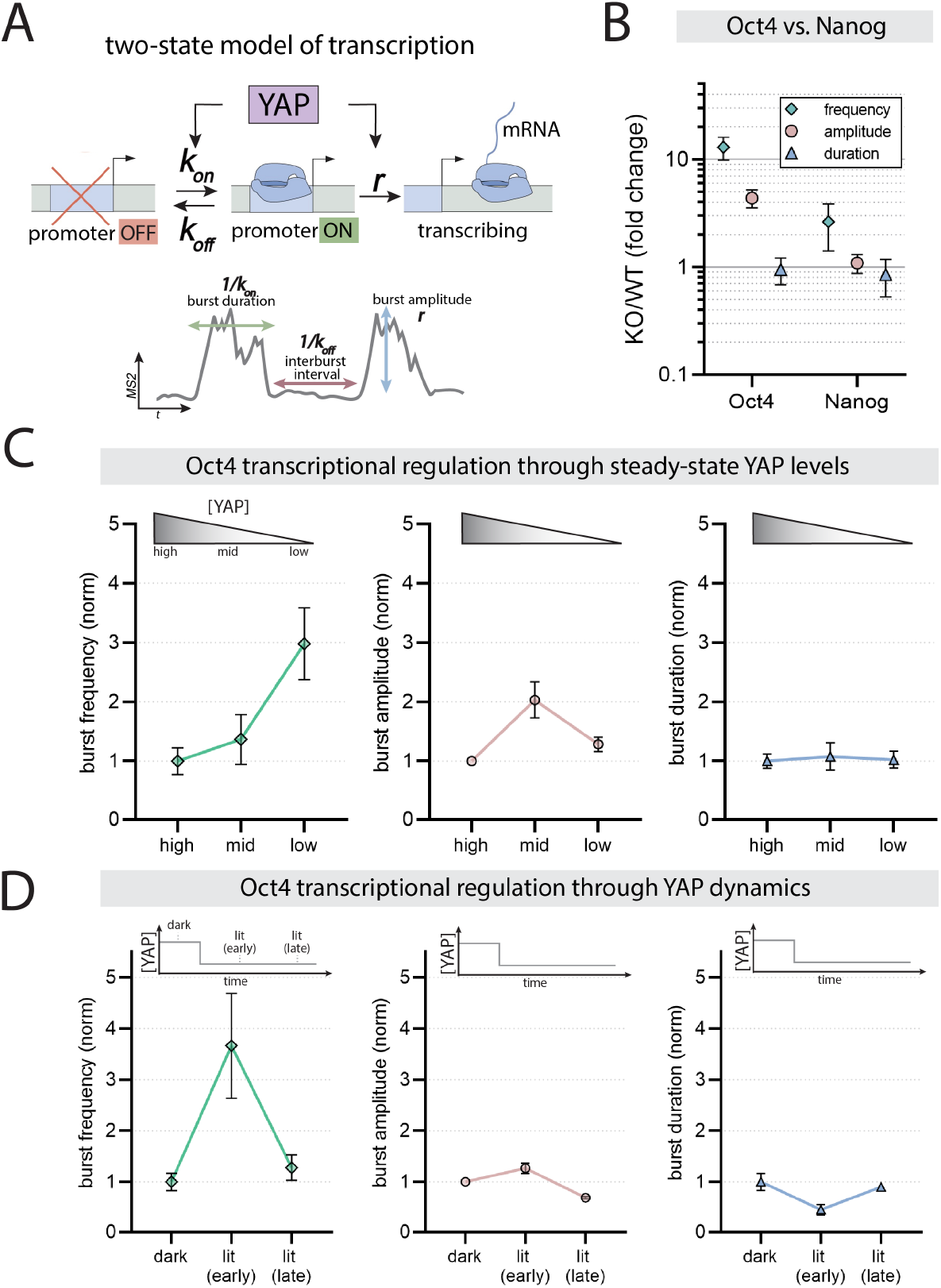
YAP regulates Oct4 transcription through modulation of burst frequency. A) Top: Two-state model of transcriptional regulation. We used this simple model to investigate where YAP interfaces with the transcription cycle to regulate downstream effectors with rate constants kon for activation of the promoter, koff for inactivation of the promoter, and r for transcriptional initiation. Bottom: The rate constants determine transcription burst duration, inter-burst interval, and burst amplitude. B) Compound-state Hidden Markov Model-based inference of transcription burst parameters (frequency, duration, amplitude) for Oct4 and Nanog in YAP KO cells, normalized to WT. p values comparing KO vs WT: frequency (Oct4: P< 0.002; Nanog: P=0.12), amplitude (Oct4: P<0.002; Nanog: P=0.4) and duration (Oct4: P=0.47; Nanog P=0.48). C,D) Transcription burst parameters inference results for Oct4 in response to titration of steady-state YAP levels (C) or upon acute nuclear YAP export (D). High, mid and low YAP levels relate to indicated YAP concentration regimes in Fig.4C. Dark, lit (early) and lit (late) relate to phases indicated as dark, response and adaptation in Fig.4D, respectively. For (C), p values comparing high to mid or high to low YAP levels: frequency (mid: P=0.21; low: P<0.002), amplitude (mid: P<0.003; low: P=0.007), duration (mid: P=0.41; low: P=0.44). For (D), p values comparing lit (early) vs dark or lit (late) vs dark: frequency (lit early: P=0.003; lit late: P=0.29), amplitude (lit early: P=0.1; lit late: P=9.3*10-5), duration (lit early: P=0.09; lit late: P=0.39).

## Discussion

YAP is a master regulator of cellular decision making in many developmental contexts. How cells achieve signaling specificity to map the right input to the right gene regulatory programs through the single node of YAP has remained an open question. Here we investigate how cells decode YAP levels and timing by using an optogenetic approach to directly manipulate YAP. Our work reveals both a concentration-dependent and dynamic signaling roles of YAP in controlling the pluripotency factors Oct4 and Nanog (Fig. 6). Because Oct4 and Nanog have different thresholds of YAP for repression, cells can titrate YAP doses for either joint or mutually exclusive regulation of Oct4 and Nanog (Fig. 6A). We further demonstrate that cells are also sensitive to the timing of YAP activation. Oscillatory dynamic YAP inputs more efficiently induce Oct4 and Nanog than do sustained inputs (Fig. 6B). By applying a range of synthetic frequencies (min-hour time scale) that bracket endogenous dynamics in mESCs, we find that natural dynamics fall into the optimum frequency decoding range suggesting they represent physiological communication codes. Together, our results identify new cellular signaling strategies to achieve the complex regulatory requirements of Oct4 and Nanog (Fig. 6C). While low steady-state YAP levels could provide a means to induce both Oct4 and Nanog for pluripotency maintenance, the offset threshold systems or dynamic inputs could preferentially induce Oct4 for mesendoderm specification. Similarly, high YAP levels may permit other lineage specification programs to act through removal of Oct4 and Nanog. While YAP has been established as a central determinant of development, its exact regulatory function has remained elusive due to conflicting reports on its requirement for pluripotency maintenance and differentiation (Chung et al., 2016; Lian et al., 2010; Sun et al., 2020). These studies have relied on genetic YAP perturbation strategies and single time point analysis of YAP levels. Our identified concentration-dependent and dynamic decoding modes were made possible by our direct manipulation and measurement of YAP in living cells in conjunction with dynamic readouts of YAP target transcriptional activation. Furthermore, by comparing naïve (2i+LIF) and differentiating cells, we demonstrate important differences in mESC signaling competence to YAP inputs. Under naïve culture conditions, Oct4 and Nanog are insensitive to a wide range of steady-state YAP concentrations, suggesting that the remodeling of the pluripotency network during pluripotency exit imparts YAP sensitivity. Our results not only address the long-standing debate on the function of YAP but also provide new strategies to rationally correct YAP signaling in disease states through targeting YAP dynamics.

**Fig. 6.**
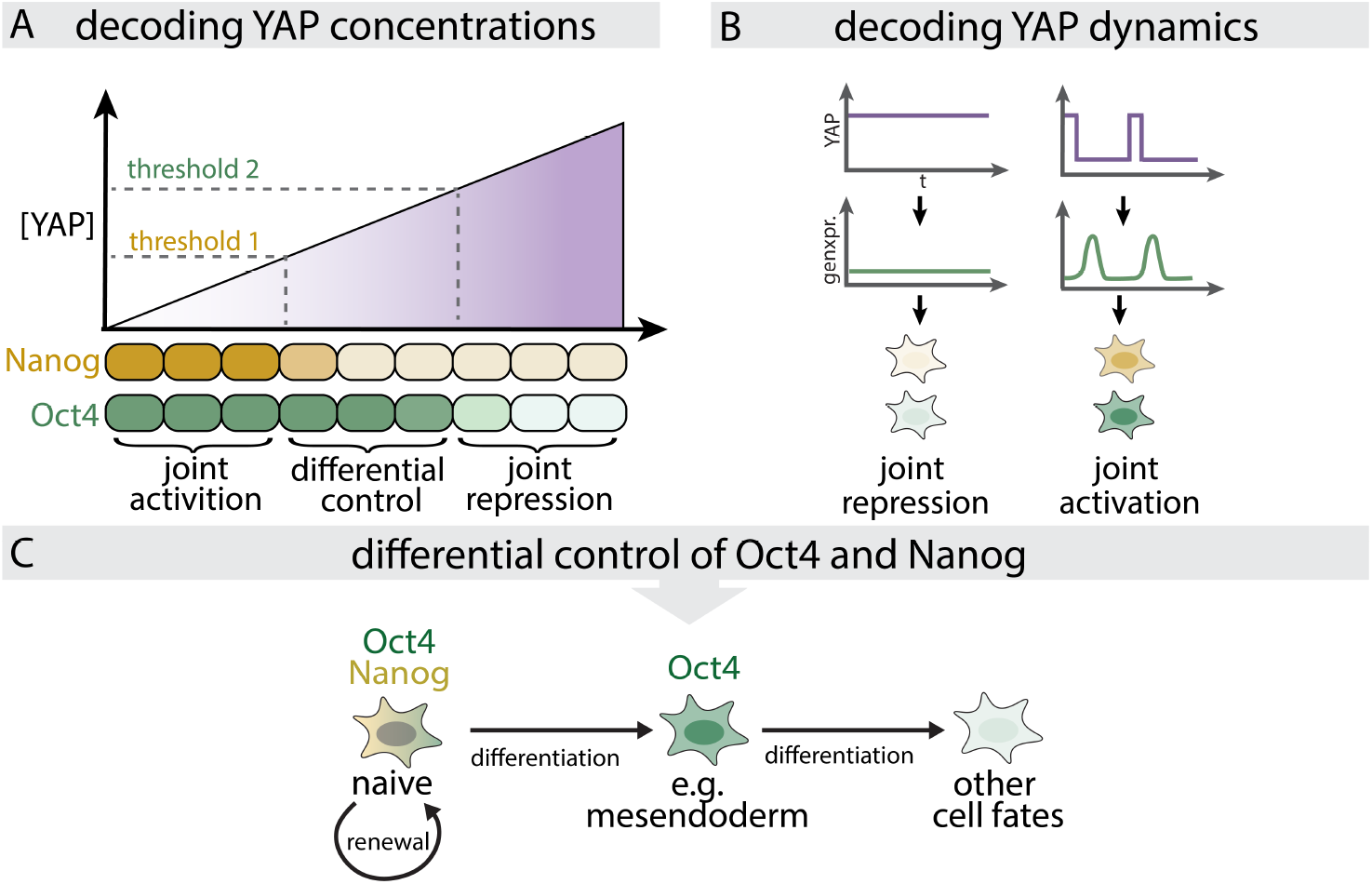
Regulation of the pluripotency factors Oct4 and Nanog through concentration-dependent and dynamic modes of YAP signaling. YAP concentrations and dynamics establish differential control strategies for the pluripotency factors Oct4 and Nanog. A) Decoding YAP steady-state concentrations through different sensitivity thresholds establish windows for the joint activation (low YAP) and repression (high YAP) or differential control (intermediate YAP) of Oct4 and Nanog. B) Decoding YAP dynamics through an adaptive change sensor for the joint control of Oct4 and Nanog expression magnitude. C) The different decoding modes depicted in A and B provide a means for the complex regulatory requirements of Oct4 and Nanog for naïve mESCs and lineage commitment.

Why are YAP effectors like Oct4 and Nanog sensitive to a particular frequency of pulsatile YAP dynamics? We identify an adaptive circuit that generates a transient burst of Oct4 transcription following an acute drop of YAP concentrations. The adaptive response operates on the hour time scale, shows a significant delay between YAP change and transcription onset (45min), and operates by transient modulation of transcription burst frequency that resets to baseline under continuous YAP export. How these response characteristics relate to the underlying molecular machinery acting at the Oct4 locus, what genetic elements the YAP-locus interaction involves (e.g. enhancer, promoter), and how the transcriptional response is shut off under continuous YAP export remain important open questions.

The use of signaling dynamics to encode information is widespread in biology. Many other transcriptional regulators such as p53, NFkB or Erk make use of dynamic communication codes to pair upstream inputs to physiologically-relevant responses (Nelson, 2004; Purvis et al., 2012; Toettcher et al., 2013). What is the advantage of dynamic modes as compared to steady-state cell signaling? Cells live in noisy environments where steady-state concentrations are subject to significant fluctuations. Dynamic readouts, such as the change sensor identified for Oct4, provide robustness to these fluctuations and are only engaged when specific temporal patterns are induced. Similarly, behavioral coordination is crucial for proper formation of developmental shape and pattern, and temporal signals provide a means to synchronize cellular decision-making. For example, Oct4-mediated induction of mesendoderm differentiation is most potently induced in G1 phase of the cell cycle (Strebinger et al., 2019). Dynamic YAP signaling could synchronize Oct4 expression with the cell cycle of either individual cells or entire cell populations. Lastly, dynamic signaling decoders often act independent of absolute concentrations but make use of relative measures such as fold-changes (Alon, 2019; Goentoro et al., 2009; Shoval et al., 2010). This scale invariance can significantly extend the dynamic range of signaling systems permitting control of cell signaling in more diverse environments than fixed concentration regimes would allow.

## ACKNOWLEDGEMENTS

We thank Jeffrey Alexander for experimental help and all members of the Weiner lab for their support. The LEXY plasmid (Addgene plasmid 72655) was a gift from the Di Ventura and Eils labs. The Janelia fluor dyes were kindly provided by Luke Lavis.

## Funding

ODW was supported by National Institutes of Health grant GM118167 and the National Science Foundation Center for Cellular Construction (DBI-1548297). KM was supported by postdoctoral fellowships from the German Research Foundation (DFG; ME 5071) and the American Heart Association (20POST35180100). HGG was supported by the NIH Director’s New Innovator Award (DP2 OD024541-01), the NSF CAREER Award (1652236), an NIH R01 Award (R01GM139913), and the Koret-UC Berkeley-Tel Aviv University Initiative in Computational Biology and Bioinformatics. HGG is also a Chan Zuckerberg Biohub Investigator.

## Supplement

**Movie S1 (related to Fig. 1B,C)**

Time-lapse confocal images of endogenous YAP (SNAP-YAP) and quantification of mean nuclear YAP intensity of the outlined nucleus (magenta) show pulsatile nuclear YAP dynamics. Cells were imaged at 1.5d post directed mesoderm induction. Scale bar 10μm.

**Movie S2 (related to Fig. 1G,H)**

Time-lapse confocal images of optoYAP expressing mESCs with simultaneous light illumination shows reversible light-gated YAP export. Blue rectangle indicates illumination phases. Scale bar 20μm.

**Movie S3 (related to Fig. S3B,C)**

Maximum projection of a time-lapse confocal z-stacks of the Oct4-MS2 transcriptional reporter in WT and YAP KO mESCs shows upregulated Oct4 expression in YAP depleted cells. Transcription spots are indicated by red circles. Scale bar 10μm.

**Movie S4 (related to Fig. 4D)**

Left: Maximum projection of time-lapse confocal z-stacks of the Oct4-MS2 transcriptional reporter in optoYAP expressing cells shows an adaptive transcriptional response upon illumination-induced YAP export. The white arrow indicates the transcription site. The illumination phase is indicated by the blue rectangle. Right: Quantification of the MS2 spot shown in the movie. The vertical line in the graph indicates the onset of the illumination phaseScale bar 10μm.

**Fig. S1.**
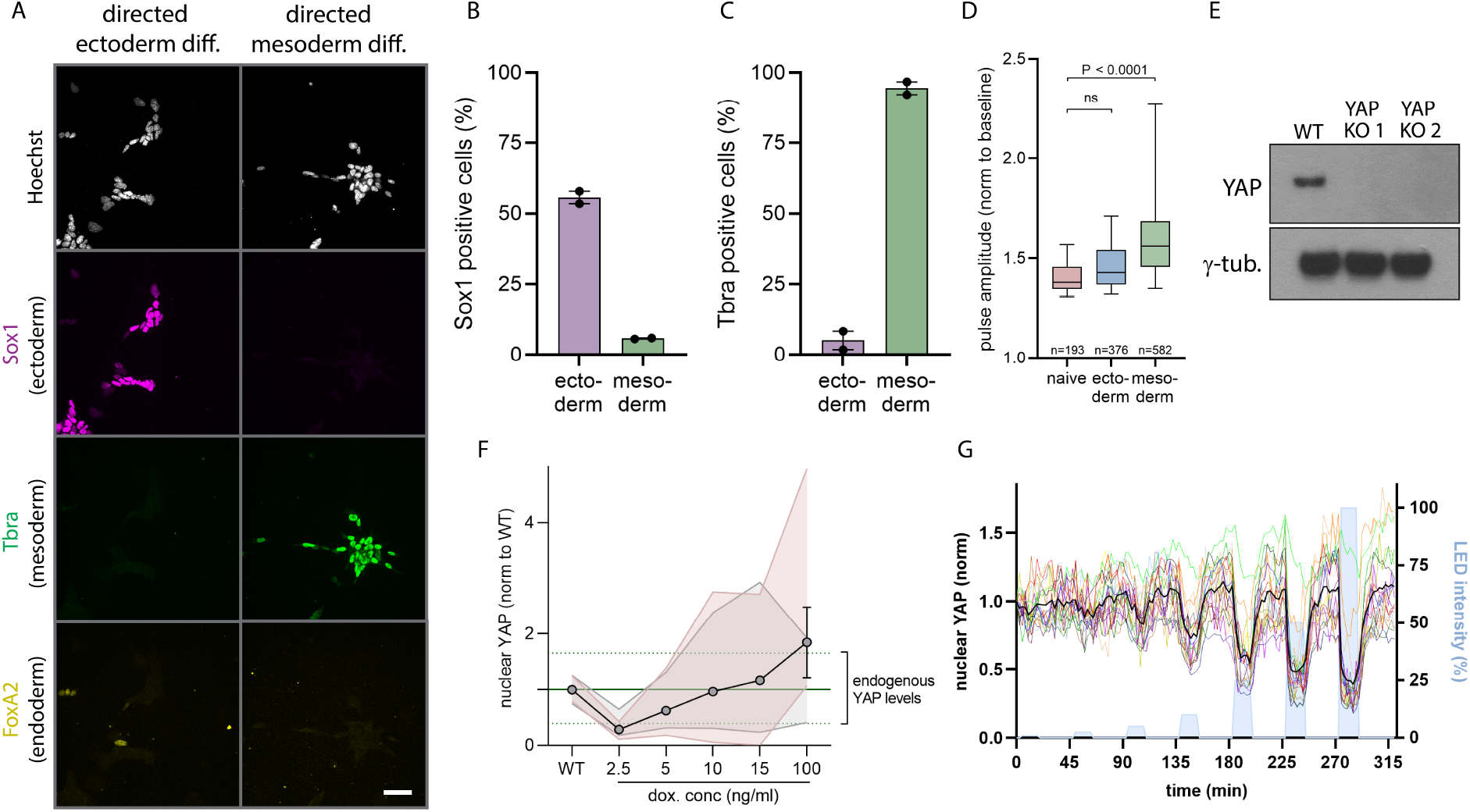
Characterization of the optoYAP tool. A) IF staining for differentiation markers Sox1 (ectoderm), Tbra (mesoderm) and FoxA2 (endoderm) at 5d post directed differentiation into the ecto- and mesoderm lineage. B, C) Quantification of Sox1 (B) and Tbra (C) positive cells upon directed differentiation from IF images as shown in (A). D) Quantification of pulse amplitude in naive mESCs versus 36h directed induction along the ectoderm or mesoderm lineages. Pulses were classified by the peak detection strategy shown in Fig. 1C. Shown are the Box and Whiskers with median and 5-95 Percentile, n as indicated, N=3. (D). p values from unpaired Student’s t test. E) Western blot detection of YAP protein levels in WT and YAP knockout (KO) mESCs shows depletion of YAP protein in two different clonal KO lines generated with different CRISPR guides. Only YAP KO 1 was used in this study. gamma-tubulin serves as loading control. F) Doxycycline induction of optoYAP in YAP KO mESCs provides access to a wide expression range bracketing the endogenous YAP levels of WT mECS. Induction was performed with indicated doxycycline concentrations. YAP levels were measured at 2d post spontaneous differentiation. The median (solid green line) + 5-95 Percentiles (dashed green lines) of the endogenous YAP level distribution are shown. OptoYAP data from two different experiments (red shading, gray shading) are shown as overlay. G) Quantification of nuclear optoYAP levels upon consecutive illumination cycles with different LED intensities demonstrates titratable nuclear YAP export. Data was quantified from microscopy time courses as shown in Fig. 1G. Illumination phases are indicated by blue shading. Colored traces represent single cells, and black solid line represents population mean.

**Fig. S2.**
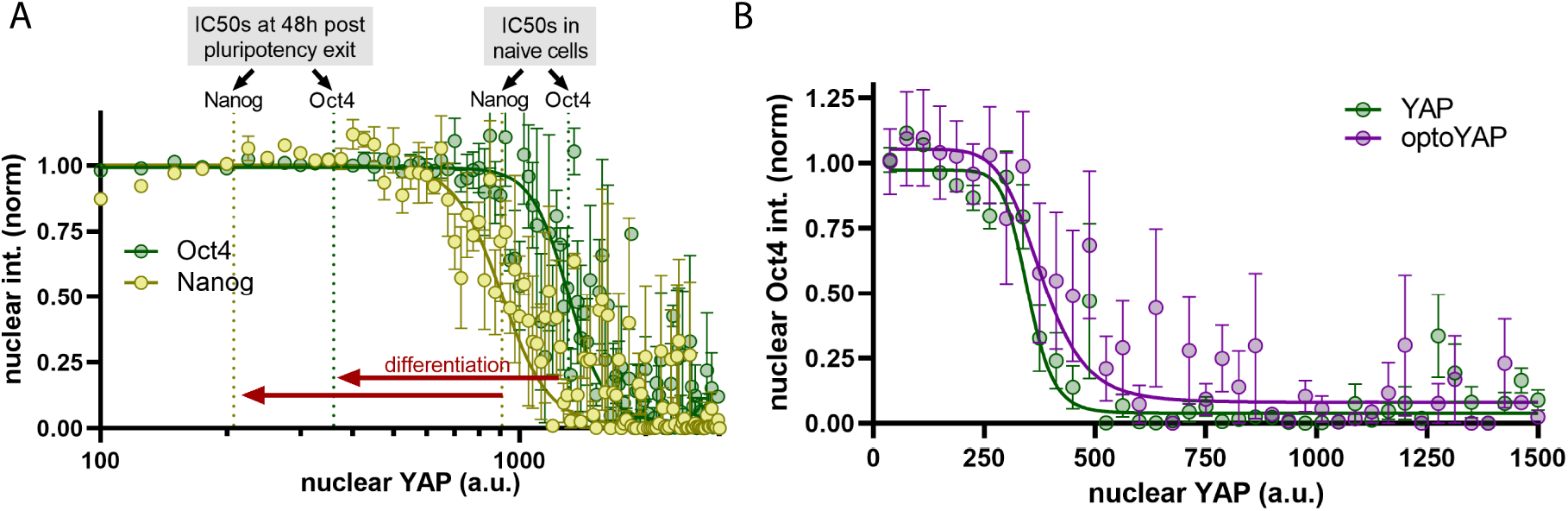
Characterization of YAP-dependent Oct4 and Nanog repression. A) Hill curve fit of nuclear Nanog and Oct4 levels as a function of nuclear YAP concentrations under naïve conditions (2i+LIF) reveals shifted repression regimes (red arrows) as compared to differentiation conditions (48h post pluripotency exit). IC50s in naïve cells and 48h post pluripotency exit for Nanog and Oct4 are indicated. The IC50s at 48h post pluripotency exit represent the data shown in Fig. 2C. Shown are mean +/-SEM, N=3. B) Comparison of the Hill curve fits of nuclear Oct4 levels as a function of nuclear YAP (green) or optoYAP (magenta) concentrations shows comparable repressive potency of both constructs. Shown are mean +/-SEM, N=4.

**Fig. S3.**
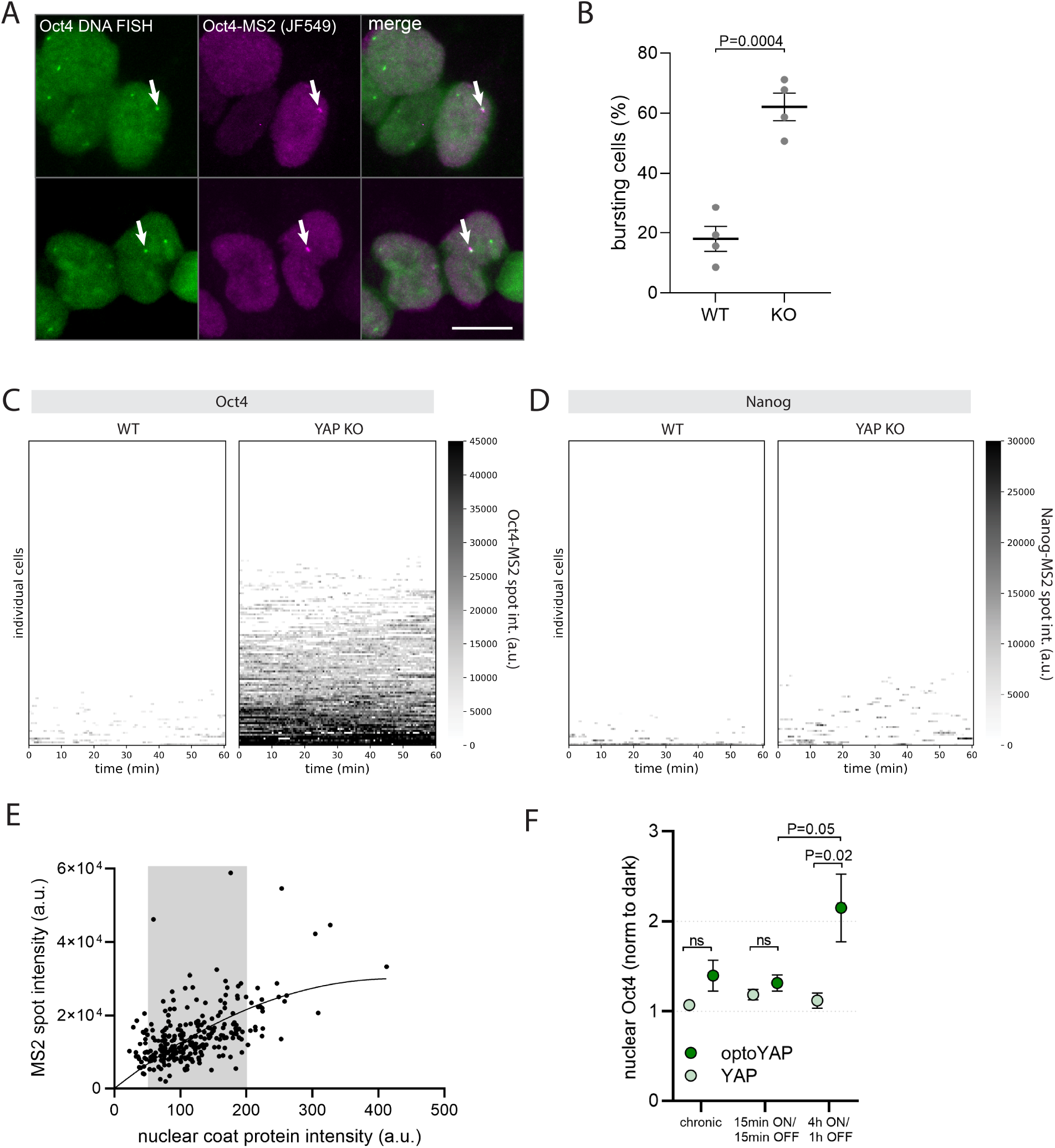
Quantification of Oct4 and Nanog transcription in YAP KO cells. A) Co-localization of Oct4-MS2 spots to the Oct4 locus visualized by Oct4 DNA FISH. Scale bar: 10 μm. B) Quantification of the percentage of Oct4-MS2 cells bursting in WT and YAP KO mESCs at 2d post pluripotency exit. Shown are mean +/-SEM, N=4. C, D) Single cell traces (y-axis) of Oct4-MS2 (C) and Nanog-MS2 (D) intensities in WT and YAP KO cells. E) Quantification and polynomial fit of the Oct4-MS2 spot intensity as a function of nuclear coat protein. Only nuclei with coat protein expression range 50-200 a.u. (grey shading) were used for quantifications shown in Fig. 4C and D. The MS2 signal was corrected for coat protein expression differences. F) Comparison of Oct4 induction on illumination with different pulsed (15min lit/15min dark or 4hlit/1h dark) or chronic light patterns shows significantly higher induction in slow YAP import/export durations (4h lit/1h dark) compared to fast or sustained ones. Culture conditions are the same as described for Fig. 3B-D. Data for the chronic and 4hON/1hOFF conditions are reproduced from Fig. 3C. Shown are mean +/-SEM, p values from unpaired Student’s t test, N=7.

## Experimental details

### mESC culture

E14 mESCs were maintained on gelatin coated dishes in 2i+LIF media, composed of a 1:1 mixture of DMEM/F12 (Thermo Fisher Scientific, 11320–033) and Neurobasal (Thermo Fisher 21103–049) supplemented with N2 supplement (Thermo Fisher 17502–048), B27 with retinoid acid (Thermo Fisher 17504–044), 0.05% BSA (Thermo Fisher 15260–037), 2 mM GlutaMax (Thermo Fisher 35050–061), 150 μM 1-thioglycerol (Sigma, M6145), 1 μM PD03259010 (Selleckchem Houston, TX, 1036), 3 μM CHIR99021 (Selleckchem S2924) and 106 U/L leukemia inhibitory factor (Peprotech, 250–02).

### Directed and spontaneous differentiation

For directed differentiation of endogenously tagged SNAP-YAP mESCs into the mesoderm lineage, cells were seeded at 10 000 cells per well on natural mouse laminin-coated 96 glass bottom plates in 2i+LIF media (see above). 12h post seeding, cells were washed three times with DMEM/0.05% BSA and cultured in differentiation media composed of a 1:1 mixture of DMEM/F12 (Thermo Fisher Scientific, 11320–033) and Neurobasal (Thermo Fisher 21103–049) supplemented with N2 supplement (Thermo Fisher 17502–048), B27 without retinoid acid (Thermo Fisher 12587010), 0.05% BSA (Thermo Fisher 15260–037), 2 mM GlutaMax (Thermo Fisher 35050–061), 150 μM 1-thioglycerol (Sigma, M6145), 3μM CHIR99021 (Selleckchem S2924). Media was changed daily. Cells were imaged at 1.5d post differentiation or fixed at day 5 for IF stainings.

For directed differentiation of endogenously tagged SNAP-YAP mESCs into the ectoderm lineage, cells were cultured in 2i+LIF SFES media in presence of 15% ES-qualified FBS for 24h. Cells were seeded at 5 000 cells per well on natural mouse laminin-coated 96 glass bottom plate in 2i+LIF media (see above) supplemented with 15% ES-qualified FBS. 12h post seeding, cells were washed three times with DMEM/0.05% BSA and cultured in differentiation media composed of a 1:1 mixture of DMEM/F12 (Thermo Fisher Scientific, 11320–033) and Neurobasal (Thermo Fisher 21103–049) supplemented with N2 supplement (Thermo Fisher 17502–048), B27 without retinoid acid (Thermo Fisher 12587010), 0.05% BSA (Thermo Fisher 15260–037), 2 mM GlutaMax (Thermo Fisher 35050–061), 150 μM 1-thioglycerol (Sigma, M6145). 24h post differentiation start, 1uM retinoic acid was added to the media. Media was changed daily. Cells were imaged at 1.5d post differentiation or fixed at day 5 for IF stainings.

For spontaneous differentiation experiments in Fig. 2B-C, Fig. 3B-D, Fig. S1F, Fig. S2A,B and Fig. S3F, cells were seeded at 15 000 cells per well on natural mouse laminin-coated 96-well glass bottom dishes in 2i+LIF media (supplemented with 1% ES-qualified FBS and 100ng/ml doxycycline) prior to spontaneous differentiation start. For live imaging experiments in Fig. 4B-D and Fig. S3B-E, cells were directly seeded at 15 000 cells per well in spontaneous differentiation media. Media was exchanged every 24h. Spontaneous differentiation media was composed of DMEM high glucose, 15% ES-qualified FBS, 2mM L-Glutamine, 0.1mM non-essential amino acids, 150 uM thioglycerol and 100ng/ml doxycycline for induction of optoYAP and YAP expression. For doxycycline dose titration experiments (Fig. S1F), cells were seeded at 1000 cells per well in a 96-well glass bottom dish in 2i+LIF media (supplemented with 1% ES-qualified FBS), containing doxycycline concentrations as indicated in Fig. S1F. 12h post seeding, cells were differentiated in spontaneous differentiation media in presence of the respective doxycycline concentrations. Cells were fixed 2d post differentiation start.

### CRISPR editing and reporter cell line generation

For CRISPR editing, we used the sgRNA/Cas9 dual expression plasmid pX330 (Addgene Plasmid 42230) and inserted sgRNA coding sequences targeting the YAP, Oct4 and Nanog locus. Homology arm sequences for generation of knockin donor vectors were amplified from E14 cDNA. pX330 and knockin donor plasmids were introduced into mESCs by electroporation using the Neon Transfection System (Thermo Fisher Scientific). Cells were transfected with 400ng pX330 plasmid and 600ng donor plasmid per 150 000 cells and electroporated with the following settings: 1400V, 10 ms pulse width, three pulses. Cells were recovered for 2 days in 2i+LIF media prior to clonal isolation or drug selection. For the generation of the YAP KO line, the YAP start sequence was targeted using the guide sequence 5’-CGGCTGTTGCGCGGGCTCCA-3’. The isolated clone used throughout this work had a 1bp insertion in the second codon of the endogenous YAP sequence, resulting in a frame shift and premature stop codon. For the generation of the endogenous SNAP-YAP reporter line in the WT mESC background, the same guide as for the YAP KO line was used (guide: 5’-CGGCTGTTGCGCGGGCTCCA-3’) and co-transfected with a SNAP-YAP donor plasmid for homologous recombination. We constructed a donor plasmid that inserted a SNAP-tag sequence upstream of the start of the YAP coding region using flanking homology arms of 800bp. For knockin of the 24xMS2 array into the Oct4 locus in WT or YAP KO mESCs, we constructed a donor plasmid that inserted a 24xMS2 cassette, followed by a start codon, puromycin coding sequence and P2A sequence directly upstream of the first exon of the Oct4 locus using homology arms of 500bp. The sgRNA used for knockin of this cassette targeted the Oct4 5’UTR (5’-TTTCCACCAGGCCCCCGGCT-3’). The isolated clones (WT or YAP KO background) harbor a single allele with the MS2 cassette. For knockin of the 24xMS2 array into the Nanog locus in the 5’ UTR in WT and YAP KO mESCs, we constructed a donor plasmid that inserted a P2A sequence followed by a puromycin coding sequence upstream of the endogenous Nanog stop codon followed by a 24x MS2 array. We used Nanog homology arms of 800bp. The sgRNA used for knockin of this cassette targeted the Nanog 3’UTR (5’-GTATGAGACTTACGCAACATC-3’). The isolated clones (WT or YAP KO background) harbor a single allele with the MS2 cassette.

For all MS2 reporter lines, we introduced the MS2 coat protein fused to two copies of the Halo-tag using the ePiggyBac transposase knockin system to generate stable lines. The MS2 coat protein was also used as nuclear marker for the endogenous SNAP-YAP cell line. All MS2 coat protein expressing lines were sorted for optimal MS2 coat protein expression by FACS.

### Cloning of doxycycline inducible optoYAP and YAP constructs

The YAP sequence used for all expression constructs was amplified from E14 mESC cDNA and represents the mouse isoform that lacks exon 6. The iLEXYi sequence (Kögler et al. 2021; Niopek et al. 2016) was generated by point mutation (V416I) of the LEXY sequence from the NLS-mCherry-LEXY plasmid (addgene 72655). An NLS-SNAP-iLEXYi cassette was fused to the N-terminus of YAP and expressed under a doxycycline inducible cassette. As non-light responsive controls, we expressed the same construct but lacking the iLEXYi sequence. The non-light responsive YAP control for the Oct4-MS2 live imaging experiments contained the additional NLS at the N-terminus of the SNAP-tag (Fig. 4C,D). The non-light responsive YAP control used in Fig. 2B,C, Fig.3 B-D, Fig. S2 A,B and Fig. S3F lacked the additional NLS sequence.

### Transient transfection of mESCs with optoYAP and YAP vectors

For all optogenetic experiments, YAP KO mESCs were transiently transfected with doxycycline inducible optoYAP and non-light responsive YAP vectors. To this end, 5*106 mESCs were electroporated with 6.6 μg plasmid using the Neon Transfection System (Thermo Fisher Scientific). Neon settings for the electroporation were as follows: 1400V, 10 ms pulse width, three pulses. Following electroporation, cells were seeded in 2i+LIF media supplemented with 1% ES-qualified FBS and 100ng/ml doxycycline. 24h post electroporation, cells were stained for 30min with SNAP-tag ligand JF646 at 10nM in 2i+LIF media (supplemented with 1% ES-qualified FBS and 100ng/ml doxycycline), washed twice and incubated for 1h in 2i+LIF media (supplemented with 1% ES-qualified FBS and 100ng/ml doxycycline) prior FACS sorting for positive cells. Positive cells were seeded for spontaneous differentiations as described in the “Directed and spontaneous differentiation” section.

### Induction of light-gated YAP dynamics and readout of Oct4 and Nanog protein levels

For quantification of Oct4 and Nanog protein levels upon exposure to different light illumination patterns, cells were transiently transfected with doxycycline inducible optoYAP and YAP constructs (see above), sorted for positive cells by FACS and seeded on PDMS subtrates with 64kPA stiffness (Advanced Biomatrix, 5261) in 96-well glass bottom plates. Cells were seeded at 15 000 cells per well in 2i+LIF media supplemented with 1% FBS and 100ng/ml doxycycline. 12h post seeding, cells were washed three time with DMEM/0.05% BSA and cultured in spontaneous differentiation media (see above). 24h post differentiation start, cells were exposed to 470nm light illumination patterns in a tissue culture incubator (37C, 5% CO2) for 12h using the optoPlate-96 (Bugaj and Lim 2019) or kept in the dark. For all illumination conditions, cells expressing non-light responsive SNAP-YAP served as controls. 11h post illumination start, cells were stained with 10nM SNAP-tag ligand JF646 for 30min under continuous light illumination. Following the 12h illumination phase, cells were recovered for 20min in the dark to reimport YAP. This allowed us to bin cells based on their optoYAP and YAP expression levels for analysis. Cells were fixed in the dark with 4% PFA for 30min and washed twice with PBS prior to IF staining for Oct4 and Nanog (see section “IF staining of mESCs”).

### IF staining of mESCs

Cells were fixed with 4% PFA (EM-grade) for 30min in the dark and washed twice with PBS. Cells were permeabilized with 0.05% TritonX-100/0.075% SDS for 20min and blocked with 10% normal goat serum (for staining with Oct4 and Nanog) or 5% BSA (for staining with Sox1, Tbra and FoxA2) for 1h. Cells were incubated with primary antibody against Sox1 (Cell Signaling Technology, 4194S), Tbra (RD Systems, AF2085), FoxA2 (Santa Cruz, sc-374376), Oct4 (Santa Cruz, sc-5279) or Nanog (Cell Signaling Technology, 8822S) in blocking buffer for 2h at room temperature, washed three times with 0.01% TritonX-100 and incubated with Alexa-488, -568 or -647 conjugated secondary antibody (1:1000, Thermo Fisher Scientific) and NucBlue (Thermo Fisher Scientific, R37605) in blocking buffer for 1h at room temperature. Cells were washed 3x 15min with 0.01% TritonX-100 and incubated in PBS for imaging.

### Imaging of IF stained samples

IF stainings were imaged on a Nikon Eclipse Ti inverted confocal microscope equipped with a CSU-W1 Yokogawa spinning disk (Andor), an iXon Ultra EMCCD camera (1024 × 1024 FOV, Andor), and 405, 440, 488, and 561-nm laser lines using a 40X Plan Apo TIRF 0.95 NA air objective (Nikon). Wells were imaged as non-overlapping 10 × 10 or 8 × 8 image grids. Pixel size was 0.325μm.

### Live imaging of endogenous YAP dynamics

For live imaging of endogenous SNAP-YAP, cells were seeded on 96-well glass bottom dishes and directed into the mesoderm and ectoderm lineages as described (see section “Directed and spontaneous differentiation“). For the naïve condition, cells were seeded at 10 000 cells per well on a glass bottom dish in 2i+LIF media and kept in 2i+LIF media throughout the experiment. The SNAP-YAP mESCs also express the MS2 coat protein (see section “CRISPR editing and reporter cell line generation”) as nuclear marker. To visualize YAP and the nuclear marker, cells were stained with SNAP-tag ligand JF646 and Halo-tag ligand JF549 (Grimm et al. 2015) at 10nM each in the respective culture media for 30min. Cells were washed three times with their respective culture media and incubated for 1h before imaging. Cells were imaged on an environmentally controlled (37C, 7% CO2) Nikon Eclipse Ti inverted confocal microscope with the same specifications as described for the imaging of the IF staining (see above). Cells were imaged with a 60x Apo TIRF 1.49 NA oil objective (Nikon) at 10 min interval for a total of 16h. Pixel size was 0.216μm.

### Live imaging of transcription and optogenetic control of YAP

For characterizing the optoYAP tool (Fig. 1H, Fig. S1G), optoYAP expressing mESCs were stained with SNAP-tag ligand JF646 and imaged on an inverted confocal microscope as described for the live imaging of endogenous YAP dynamics (see above). Cells were simultaneously illuminated with 470nm light from top using the optoPlate-96 (Bugaj and Lim 2019). The optoPlate-96 was mounted on the stage such that the temperature and CO2 control was maintained. Light was applied as pulses with a 1 sec ON/1sec OFF duty cycle.

For live imaging of Oct4 and Nanog transcription in WT and YAP KO cell lines harboring the MS2 reporter, as well as for simultaneous live imaging of optoYAP or YAP and Oct4-MS2 transcription, 50 μg/mL ascorbic acid and a 1:100 dilution of Prolong Live Antifade Reagent (ThermoFisher P36975) were added to the media 30min prior imaging start to reduce photobleaching. Cells were imaged on an inverted Nikon Ti confocal microscope equipped with a CSU-22 spinning disk confocal, Photometrics Evolve Delta EMCCD camera (512 × 512) and a cage incubator with CO2 and humidity control.

For experiments comparing the MS2 transcriptional activity in WT and YAP KO lines (Fig. 5B, Fig. S3 C,D), cells were imaged with a Plan Apo VC 60x/1.4 NA oil objective (pixel size 154nm). Images were acquired as z-stacks with planes spaced 200nm apart, covering a total of 8-10 um in z (= 41-51 z slices) and acquired every 30 sec.

For experiments with simultaneous optogenetic control of nuclear YAP levels and imaging of the MS2 transcriptional reporter, cells were imaged with a Plan Fluor 40x/1.3 NA oil objective (pixel size 228) using a dual band pass red and far red emission filter that blocks blue light. 470nm light for optogenetic light stimulation was applied using the optoPlate-96. The plate was mounted on top of the stage such that the temperature and CO2 control was maintained. Light was applied pulsed with a 1 sec ON/1sec OFF duty cycle. The MS2 signals was acquired as z-stack with planes spaced 250nm apart, covering a total of 17 um in z (= 71 z slices) and acquired every 1.5min. The optoYAP or YAP signals was captured as single slice in the center of the stack and acquired every 22.5 min. The optoYAP and YAP control cells were imaged in the same run in adjacent wells. For all MS2 and optoYAP imaging experiments, the laser output was manually adjusted for every experiment to quantitatively compare optoYAP, YAP and MS2 intensities between experiments.

### Denoising of microscopy images

Images from time lapse microscopy of the MS2 reporter and endogenous YAP were denoised using NDSafir (Kervrann and Boulanger 2006; Carlton et al. 2010).

### Quantification of endogenous YAP levels from live imaging time courses

Denoised images were background subtracted using a dark field image and flat field corrected. Nuclei were segmented based on the nuclear marker (MS2 coat protein) using the StarDist detector (Schmidt et al. 2018) and tracked using the overlap tracker in the Fiji Trackmate plugin (Ershov et al. 2022). Small objects were filtered out and only tracks with track length > 10h were kept. Trackmate quantifications of the nuclear median YAP intensity were imported into Python (version 3.8.5) for peak detection. Cells with significant changes of the nuclear marker (see section “live imaging of endogenous YAP dynamics”) were excluded as this indicated tracking error or mitotic cells. The first and last three frames of each movie were additionally removed to ensure cells were not entering or exiting mitosis. Each trace was normalized to the average trace computed from all nuclei of a movie, to correct for bleaching and fluorescent drift. The corrected single cell traces were then smoothed using a rolling mean and peak detection was performed using the find_peaks function from the signal module of the package. This function finds all local maxima by simple comparison of neighboring values. From this, we computed the percentage of cells showing YAP pulses as well as the pulse duration and amplitude.

### Quantification of nuclear optoYAP levels

The mean nuclear optoYAP intensities (Fig. 1H and Fig. S1G) were manually quantified from time lapse movies by drawing a rectangle in the nucleus using Fiji (Schindelin et al. 2012).

### Quantification of Nanog and Oct4 from IF stainings

Nuclei were segmented based on the NucBlue (Hoechst) staining using the Fiji StarDist plugin (Schmidt et al. 2018). Small objects were filtered out and the mean nuclear Oct4, Nanog and optoYAP or YAP intensities were quantified for each nucleus. For establishing the Hill curves (Fig. 2C), the optoYAP and YAP intensities were scaled to compare between experiments. To this end, we plotted the Nanog and Oct4 intensities as a function of YAP, computed the IC50s from Hill curve fits and used the center of the Nanog and Oct4 IC50s to scale experiments. Following scaling of the optoYAP intensities, the Oct4 and Nanog signal was min-max normalized using the bottom and top plateau of each Hill curve as min and max respectively.

For quantification of the mean nuclear Oct4 and Nanog intensities upon light-gated oscillatory YAP inputs (Fig. 3), we binned the nuclei according to their optoYAP or YAP expression levels. To this end, nuclei from the dark control well were used to plot the mean nuclear Oct4 intensity as a function of nuclear optoYAP levels, to fit a Hill curve. From the fit, the IC99 was computed and used as threshold to classify cells into high YAP or optoYAP expressors (>IC99). Assuming that Oct4 and Nanog induction is most pronounced in repressed cells, only high YAP expressing cells were considered for the quantification of the median nuclear Oct4 and Nanog intensities in Fig. 3.

### Quantification of Sox1 and Tbra positive cells for directed differentiations

Nuclei were segmented using the NucBlue channel as described for the quantification for nuclear Nanog and Oct4 from IF stainings (see above). Segmentation masks were used to uquantify median nuclear Sox1 and Tbra intensities. To establish cut-off values for identification of Sox1 and Tbra positive cells, we used the the 95 percentile of the negative control as cutoff to distinguish negative and positive cells. For example, the 95 percentile of the nuclear Sox1 intensities of the directed mesoderm condition was defined as the threshold for ectoderm (=Sox1) positive cells. The percentage of positive cells was quantified to determine the efficiency of the directed lineage differentiations.

### Quantification of MS2 spots from time lapse movies

3D time-lapse images were converted into 2D images by maximum Z projection. The AI segmentation algorithm form the NIS.ai suite of the NIS-Elements software (Nikon) was used for initial MS2 spot detection and nuclear segmentations from these projections. The segmented images were fed into a Python analysis pipeline to quantify the intensity of the MS2 spot and the nuclear YAP intensities. First, the trackpy Python package was used to track individual nuclei and quantify the mean nuclear MS2 coat protein and optoYAP or YAP intensities. Nuclei that touched the image border or were dividing were excluded. Next, the MS2 spots identified by the Nikon AI were verified by a 2D gaussian fit on the maximum projection, followed by a 1D gaussian fit along the z-dimension of the image stack. Finally, the sum pixel intensities within the identified spots was quantified from the z-stack. Spot intensities were corrected for cell-to-cell differences in MS2 coat protein expression levels. To this end, the average spot intensity per nucleus was plotted against the MS2 coat protein levels (see Fig. S3E) and a second order polynomial was fit to the data. Only nuclei with non-saturating MS2 coat protein levels (50-200.a.u., see Fig. S3E) were used for quantifications of transcriptional activity. MS2 spot intensities were corrected for MS2 coat protein expression levels.

### Statistics

Graphpad was used for statistical analysis (Graphpad software, Inc). Details can be found in the legend of each figure. N represents number of independent biological replicates.

## Burst parameter inference methods

### cpHMM

The burst parameter trends shown in Figure 5 were obtained using cpHMM, a computational method that employs compound-state Hidden Markov Models to infer promoter state dynamics and burst parameter values (frequency, duration, and amplitude) from populations of single-cell traces. See (Lammers et al. 2020) for details regarding the method’s implementation. Briefly, transcriptional traces were divided into inference groups according to either KO/WT status (Figure 5B), average nuclear YAP concentration (Figure 5C), or according to time relative to optogenetic perturbation (Figure 5D). Parameter estimates for each inference group were estimated by taking the average across no fewer than 16 separate bootstrap samples. Each bootstrap sample contained at least 1000 time points. Outlier bootstrap results were excluded using MAtlab’s built-in “rmoutliers” function, which defines outliers as any value that is more than three scaled median absolute deviations from the population median. Inference uncertainty was estimated by taking the standard deviation across these bootstrap replicates. We used a model with two burst states (OFF and ON), as illustrated in Figure 5A.

Estimating elongation times A key input parameter for cpHMM inference is the amount of time required for RNA Polymerase molecules to transcribe the reporter gene. We estimated this quantity for our Nanog and Oct4 reporters by examining “low-to-high” transition events in our MS2 data. Specifically, we set a “high” threshold, *f*_*high*_, for each inference group, defined as the 85th percentile of all observed fluorescent spot intensities and a “low” threshold defined as *f*_*low*_ = 0.15*f*_*high*_. We use these thresholds to filter for instances in our MS2 traces where the system transitions from low (*f ≤ f*_*low*_) to high (*f ≥ f* _*high*_) fluorescence levels. We then took the average across these events to obtain an averaged low-to-high event.

Intuitively, the time required for the system to transition from low to high fluorescence captures how long it takes to change from having an empty (or nearly empty) reporter gene–low fluorescence state–to a full (or nearly full) reporter gene–high fluorescence state. Accordingly, we used Matlab’s built-in “findchangepts” function to estimate the number of time steps required to transition from basal fluorescence levels to saturating levels for each reporter gene, and defined this quantity as the elongation time. For the KO vs. WT experiments depicted in Figure 5B, which had an experimental time resolution of 30 seconds, this procedure produced elongation time estimates of 3 steps (90 seconds) and 8 steps (240 seconds) for Nanog and Oct4, respectively. For the optogenetic results depicted in Figure 5D, which had a time resolution of 90 seconds, we obtained an elongation time estimate of 3 steps (270 seconds) for Oct4.

### Estimating p-Values

We used bootstrap cpHMM inference replicates to calculate p-Values for the results shown in Figure 5B-D. To illustrate our approach, we describe the procedure in detail for the steady-state Oct4 response to YAP levels (Figure 5C). The same approach was used to calculate p-Values for the remaining results in Figure 5.

The panel of Figure 5C shows burst parameter results for Oct4 loci exposed to low, medium, and high YAP concentrations. For each burst parameter, the trend is normalized by the “high” YAP value, such that the results report on the fold difference in burst frequency, duration, across YAP levels. Consider the fold change in burst frequency for intermediate (“mid”) YAP levels. On average, the burst frequency increases by a factor of about 1.4 relative to high YAP levels. In this case then, we wish to establish how confident we can be in our finding that the fold change is significantly greater than 1.

To do this, we make use of our bootstrap replicates. Once outliers are removed, we have *N*_*high*_ inference bootstraps for the high condition and *N*_*mid*_ inference bootstraps for the intermediate condition. We then calculate the fold increase for all possible combinations of mid and high bootstrap results, leading to *N*_*f old*_ = *N*_*high*_*xN*_*mid*_ distinct bootstrap estimates of the fold increase. Our p-value is then simply defined as the fraction of these *N*_*f old*_ estimates that are found to be less than or equal to 1; i.e. the fraction of replicates for which the observed fold increase does not hold. If the trend is significant, this will occur only rarely, whereas this will occur frequently if a trend is small relative to its uncertainty. In this case, we find that about 20% of all bootstrap fold change values are less than or equal to 1. This equates to a p value of 0.2, which implies that the result is not significant at the 10%, 1%, or 0.1% levels. This same bootstrap-based approach is used to assess significance levels for all results shown in Figure 5.

## Notes

### Competing Interest Statement

The authors have declared no competing interest.

### Summary of Updates

author name corrected

